# Visualizing interactions of VDAC1 in live cells using a tetracysteine tag

**DOI:** 10.1101/2023.09.08.556841

**Authors:** Johannes Pilic, Furkan E. Oflaz, Benjamin Gottschalk, Yusuf C. Erdogan, Wolfgang F. Graier, Roland Malli

## Abstract

The voltage-dependent anion channel 1 (VDAC1) is a crucial gatekeeper in the outer mitochondrial membrane, controlling metabolic and energy homeostasis. The available methodological approaches fell short of accurate visualization of VDAC1 in living cells. To permit precise VDAC1 imaging, we used the tetracysteine (TC)-tag approach and visualized VDAC1 dynamics in living cells. TC-tagged VDAC1 had a cluster-like distribution on mitochondria. The majority of VDAC1-clusters were localized at endoplasmic reticulum (ER)-mitochondria contact sites. Notably, VDAC1 colocalized with BCL-2 Antagonist/Killer (BAK)-clusters upon apoptotic stimulation. Additionally, VDAC1 was found at mitochondrial fission sites, likely promoting mitochondrial fragmentation. These findings highlight the suitability of the TC-tag for live-cell imaging of VDAC1, shedding light on the roles of VDAC1 in multiple cellular processes.

## Introduction

Fluorescent proteins (FPs) have become an indispensable tool to tag and visualize proteins in living cells.^1^ However, the large size of FPs (∼25 kDa),^2^ can potentially interfere with the function and localization of the tagged protein. Such complications have been reported by GFP-tagging of the voltage-dependent anion channel 1 (VDAC1), a crucial transporter of metabolites and ions in the outer mitochondrial membrane.^3,4^ Correct insertion of VDAC1 into the membrane depends on the location and length of the protein tag.^5^ It has been shown that the presence of a C-terminal tag of eight or more residues leads to mistargeting of VDAC1,^3–5^ most likely by interfering with the recognition of the targeting signal by the mitochondrial import receptor Tom20.^6^ Based on this knowledge, we hypothesized that a fusion tag shorter than eight residues would allow live-cell imaging of correctly localized VDAC1.

The tetracysteine (TC)-tag, consisting of only six residues (CCPGCC), has emerged as a promising tool to visualize protein dynamics in living cells.^7^ The TC-tag becomes fluorescent upon binding of fluorogenic dyes, such as the **fl**uorescein **a**r**s**enical **h**airpin binder-**e**thane**d**i**t**hiol (FlAsH-EDT_2_) or the red-shifted analog **re**sorufin **a**r**s**enical **h**airpin binder-**e**thane**d**i**t**hiol (ReAsH-EDT_2_).^8^ Here, we aim to assess the functionality and suitability of the TC-tag as a tool to image VDAC1 in live cells. We anticipate that incorporating the TC-tag will enable the precise visualization of VDAC1 localization, trafficking, and interactions with organelles.

## Results

### Mistargeting of VDAC1 is induced by N- and C-terminal fusion of GFP

To assess whether VDAC1 can fold properly when fused to FPs with a flexible linker, we used AlphaFold2 to generate 3D structures for GFP-VDAC1 and VDAC1-GFP (Figure 1A). The 3D structures showed favorable folding patterns for both constructs, suggesting that the fusion of FPs does not compromise the quarternary structure of VDAC1.

**Figure 1.**
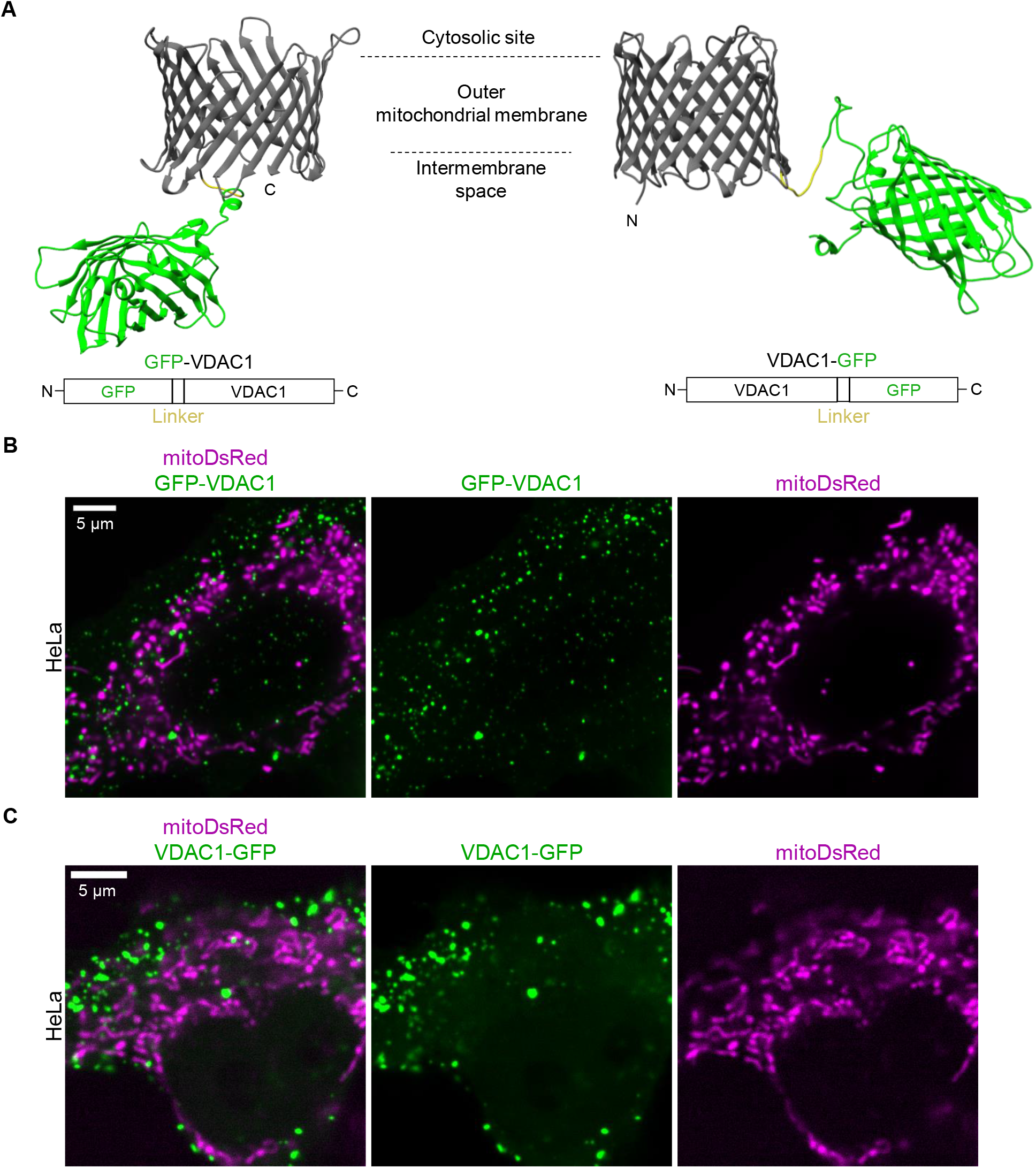
Mistargeting of VDAC1 is induced by N- and C-terminal fusion of GFP. (A) AlphaFold2-generated 3D structures and schematic representation of GFP-VDAC1 and VDAC1-GFP fusion constructs. (B) Confocal images of a HeLa cell expressing GFP-VDAC1 and mitoDsRed. (C) Confocal images of a HeLa cell expressing VDAC1-GFP and mitoDsRed.

To validate the correct targeting of these constructs, we imaged HeLa cells coexpressing GFP-VDAC1 (Figure 1B) or VDAC1-GFP (Figure 1C) with mitoDsRed, an FP targeted to the mitochondrial matrix. We observed that both constructs were predominantly aggregated and did not localize to mitochondria (Figure 1B, and Figure 1C), indicating the mistargeting of the constructs.

### Short tetracysteine-tag allows to visualize VDAC1-clusters on mitochondria

Since VDAC1 tends to mistarget when attached to a C-terminal tag of eight or more residues,^5^ we opted to fuse VDAC1 with a six-residue long tetracysteine (TC)-tag. We choose to tag the C-terminus of VDAC1, as the N-terminal domain is known to interact with numerous proteins.^9^ To visualize VDAC1-TC, we used two chemical dyes: FlAsH-EDT_2_ (Figure 2A) and the red-shifted ReAsH-EDT_2_ (Figure 2B). To determine the optimal FlAsH concentration for VDAC1 labeling, we imaged HeLa cells coexpressing VDAC1-TC and mitoDsRed with increasing FlAsH concentrations. We observed a concentration-dependent increase in green fluorescence intensity and high noise below 0.1 µM of FlAsH (Figure 2A, second row), suggesting that the optimal dye concentration lies between 0.1 and 1 µM of FlAsH. Upon closer examination, we identified two distinct labeling patterns: Transfected cells exhibited a cluster-like distribution of VDAC1 around mitochondria (Figure 2A, third row). In contrast, untransfected cells showed homogenous labeling of mitochondria by FlAsH, indicating unspecific mitochondrial accumulation of the dye (Figure 2A, fourth row). Importantly, transfection with mitoDsRed alone did not lead to a cluster-like distribution of FlAsH (Figure S1A), suggesting that the dye alone does not cluster on mitochondria.

**Figure 2.**
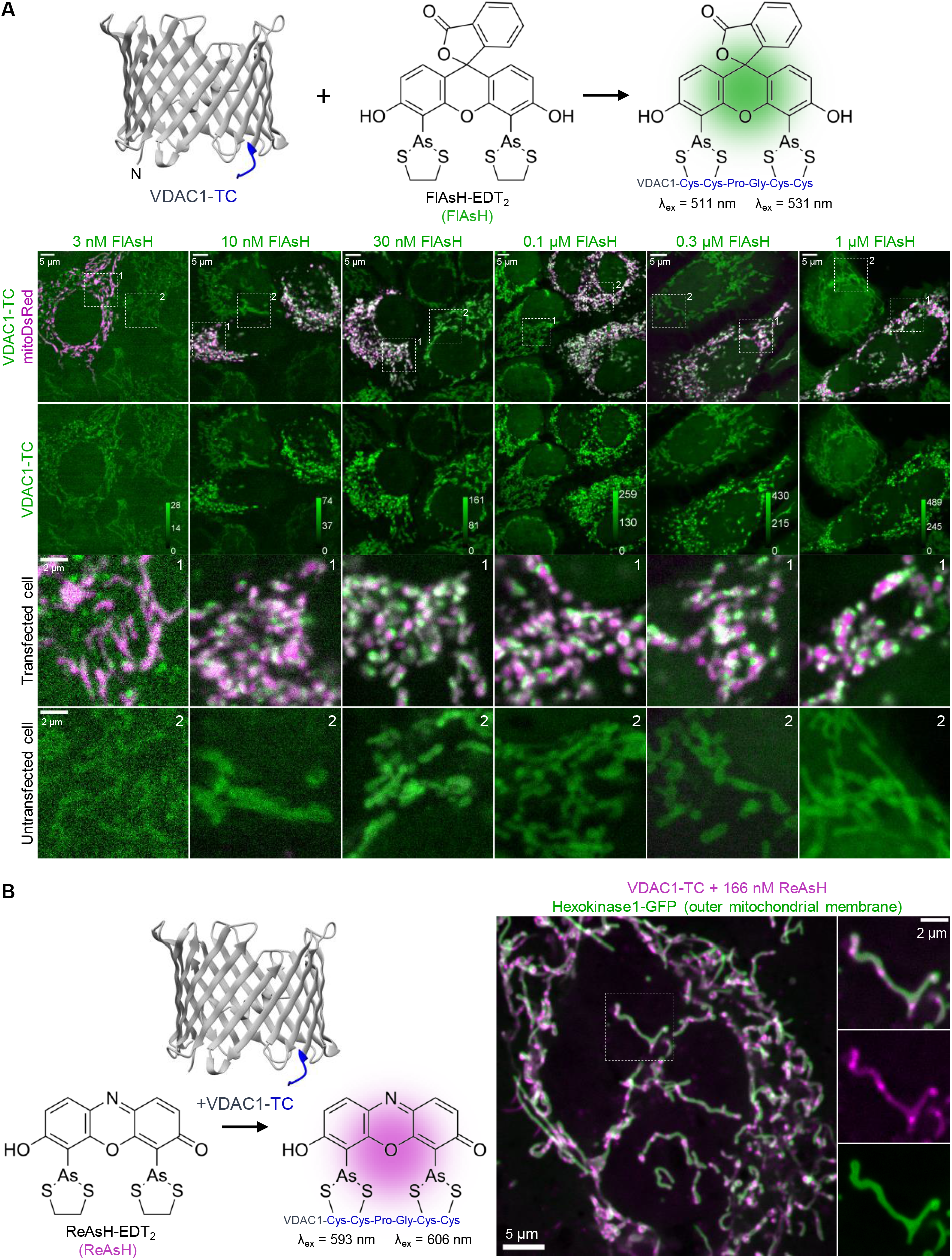
Short tetracysteine-tag allows to visualize VDAC1-clusters on mitochondria. (A) 3D structure of VDAC1-TC and chemical structure of FlAsH (top). Confocal images of HeLa cells transfected with VDAC1-TC and mitoDsRed; cells were labeled with FlAsH at increasing concentrations for 30 min (bottom). The first row of images shows an overview of the cells, and the dashed squares are magnified in the third and fourth rows. In the second row, the intensity levels of FlAsH are depicted in the calibration bar. (B) 3D structure of VDAC1-TC and chemical structure of ReAsH (left). Confocal images of a HeLa cell expressing VDAC1-TC and Hexokinase1-GFP; cells were labeled with 166 nM ReAsH for 15 min and washed with 100 µM BAL for 10 min (right).

Due to structural similarities of FlAsH with Rhodamine123 (Figure S1B), a common potentiometric mitochondrial dye, we hypothesized that unspecific binding of FlAsH to mitochondria could be reduced with carbonyl cyanide-p-trifluoromethoxyphenylhydrazone (FCCP), a mitochondrial uncoupler. Indeed, treating cells with FCCP during labeling significantly reduced the binding of FlAsH to mitochondria compared to FCCP untreated cells (Figure S1B), suggesting unspecific binding of FlAsH is caused by mitochondrial membrane potential. The classical washing buffer for the TC-labeling approach, British anti-Lewisite (BAL), also reduced the binding of FlAsH to mitochondria, but not as much as FCCP (Figure S1B).

Additionally, we tested whether overexpression of Hexokinase 1 (HK1), which has been shown to interact with VDAC1 on the outer mitochondrial membrane (OMM),^10^ affects the cluster-like distribution of VDAC1. Using HK1-GFP and ReAsH labeled VDAC1-TC we observed that VDAC1-clusters localized on the OMM (Figure 2B), suggesting that HK1 overexpression does not affect the subcellular localization of VDAC1-clusters.

### VDAC1-clusters are localized at ER-mitochondria contact sites

Since VDAC1 has been described to localize to ER-mitochondrial contact sites,^11,12^ we examined whether VDAC1-clusters colocalize with the ER in living cells. For this purpose, we imaged HeLa cells coexpressing VDAC1-TC, with the ER-marker mCh-ER3, and mitoCFP. We observed that VDAC1-clusters indeed localize to ER-mitochondrial contact sites (Figure 3A). We quantified that 63.6% of VDAC1-clusters colocalize with the ER (Figure 3B), suggesting that VDAC1 is involved in ER-mitochondria communication.

**Figure 3.**
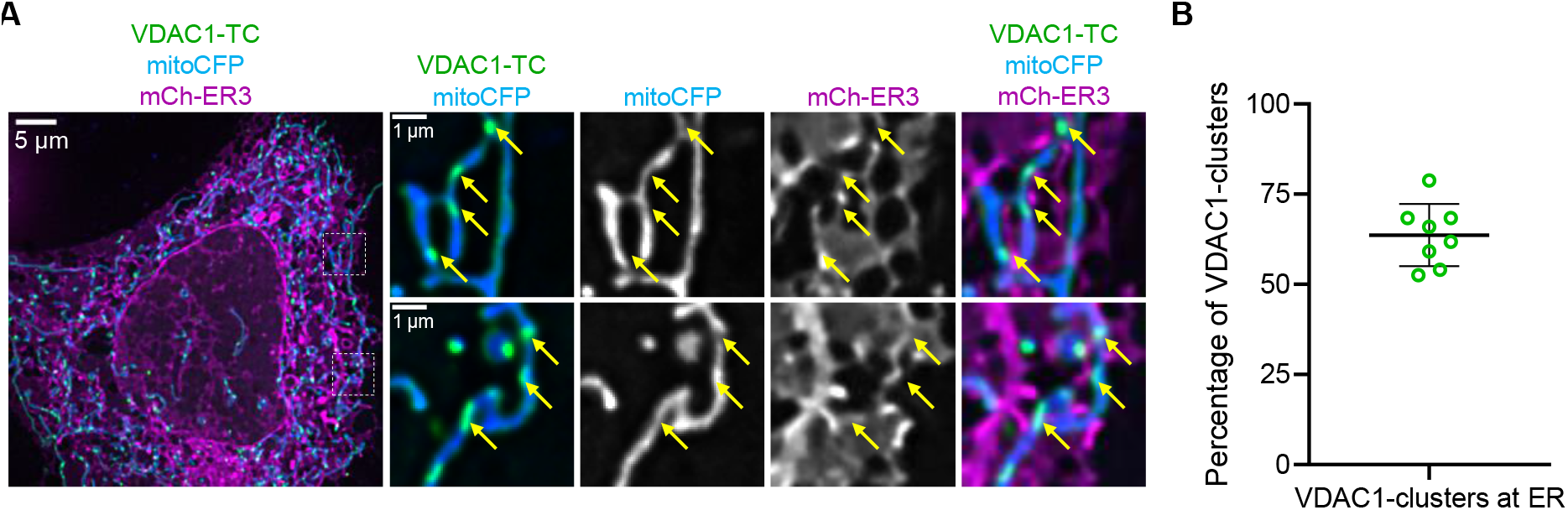
VDAC1-clusters are localized at ER-mitochondria contact sites. (A) Confocal images of a HeLa cell expressing VDAC1-TC, mitoCFP, and mCh-ER3; the cell was labeled with 1 µM FlAsH for 15 min and washed with 100 µM Bal for 10 min. The left image shows an overview of the cell, and the dashed squares are magnified on the right side. Yellow arrows point to positions of VDAC1-clusters at ER-mitochondria contact sites. (B) The graph shows the percentage of VDAC1-clusters that colocalize with the ER (n = 8).

### VDAC1 interacts with BAK-clusters that form in response to stress

VDAC1 has been suggested to interact with BCL-2 Antagonist/Killer (BAK).^13^ To explore this dynamic interaction, we imaged HeLa cells coexpressing VDAC1-TC with GFP-BAK. BAK is a proapoptotic protein residing as an inactive monomer on the OMM until triggered by cellular stress to cluster into pores. We found that glucose depletion rapidly triggered BAK-clustering (Figure 4A). Quantifying the interaction between BAK and VDAC1, we found that 39.1% of VDAC1-clusters colocalized with BAK-clusters and 23.0% of BAK-clusters colocalized with VDAC1-clusters during glucose depletion (Figure 4A). BAK-clusters with similar interaction to VDAC1 were observed upon treatment with a classical pro-apoptosis activator, staurosporine (STS) (Figure 4B). These observations provide the first evidence of VDAC1-BAK interaction during cellular stress in living cells.

**Figure 4.**
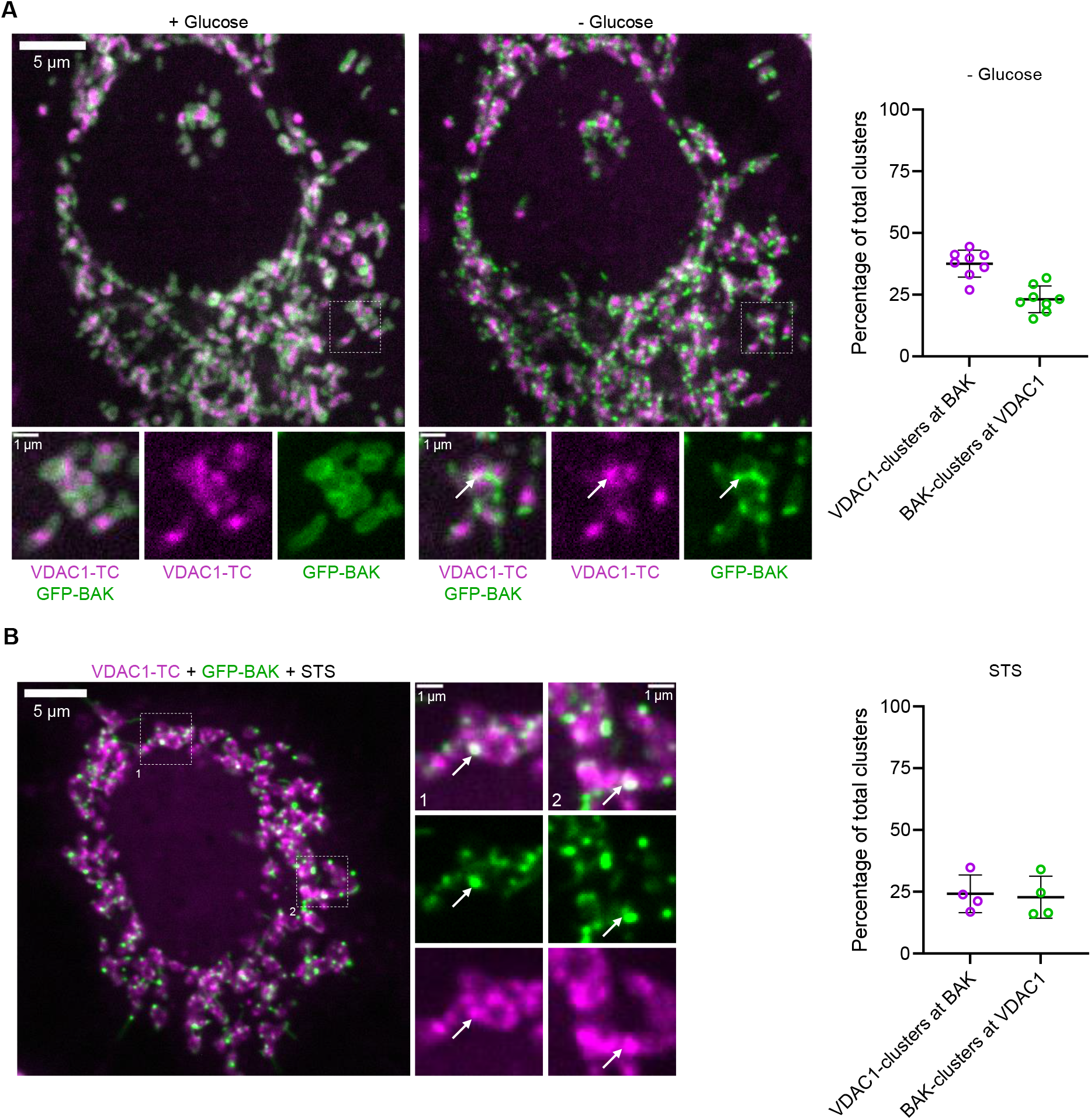
VDAC1 interacts with BAK-clusters that form in response to stress. Confocal images of a HeLa cell expressing VDAC1-TC and GFP-BAK with 10 mM glucose (left) and after 20 min of glucose depletion (middle); the cell was labeled with 333 nM ReAsH for 15 min. The first row shows an overview of the cell, and the dashed squares are magnified below. White arrows point to the contact site between VDAC1- and BAK-clusters. The graph shows the percentage of VDAC1-clusters that colocalized with BAK-clusters and the percentage of BAK-clusters that colocalized with VDAC1-clusters after 20 min of glucose depletion (right). Confocal images of a HeLa cell expressing VDAC1-TC and GFP-BAK after treatment with 10 µM STS for one hour; the cell was labeled with 333 nM ReAsH for 15 min. On the left, an overview of the cell is shown, and the dashed squares are magnified in the middle. White arrows point to the contact sites between VDAC1- and BAK-clusters. The graph shows the percentage of VDAC1-clusters that colocalized with BAK-clusters and the percentage of BAK-clusters that colocalized with VDAC1-clusters after treatment with 10 µM STS for one hour (right).

### VDAC1 overexpression promotes mitochondrial fragmentation

We noted an increased fragmentation of mitochondria in cells expressing VDAC1-TC compared to untransfected cells. Therefore, we compared the mitochondrial morphology between HeLa cells expressing mitoDsRed and HeLa cells coexpressing VDAC1-TC with mitoDsRed (Figure 5A). To assess mitochondrial morphology, we used Aspect Ratio (AR) and Form Factor (FF), which are indicators for mitochondrial swelling and branching, respectively. As evidenced by the significant reduction of AR and FF, VDAC1-TC overexpression induced mitochondrial fragmentation in HeLa cells (Figure 5A). Next, we tested whether VDAC1 is involved in mitochondrial fission events. We observed that VDAC1-clusters were localized to sites of mitochondrial fission using FCCP, a potent inducer of mitochondrial fission (Figure 5B). To test whether VDAC1 is involved in mitochondrial fission triggered by elevations in cellular Ca^2+^, we used ionomycin, a potent Ca^2+^ ionophore. Ionomycin treatment induced mitochondrial fragmentation, with VDAC1-clusters localizing to mitochondrial fission sites (Figure 5C). These observations suggest that VDAC1 plays a role in mitochondrial fission.

**Figure 5.**
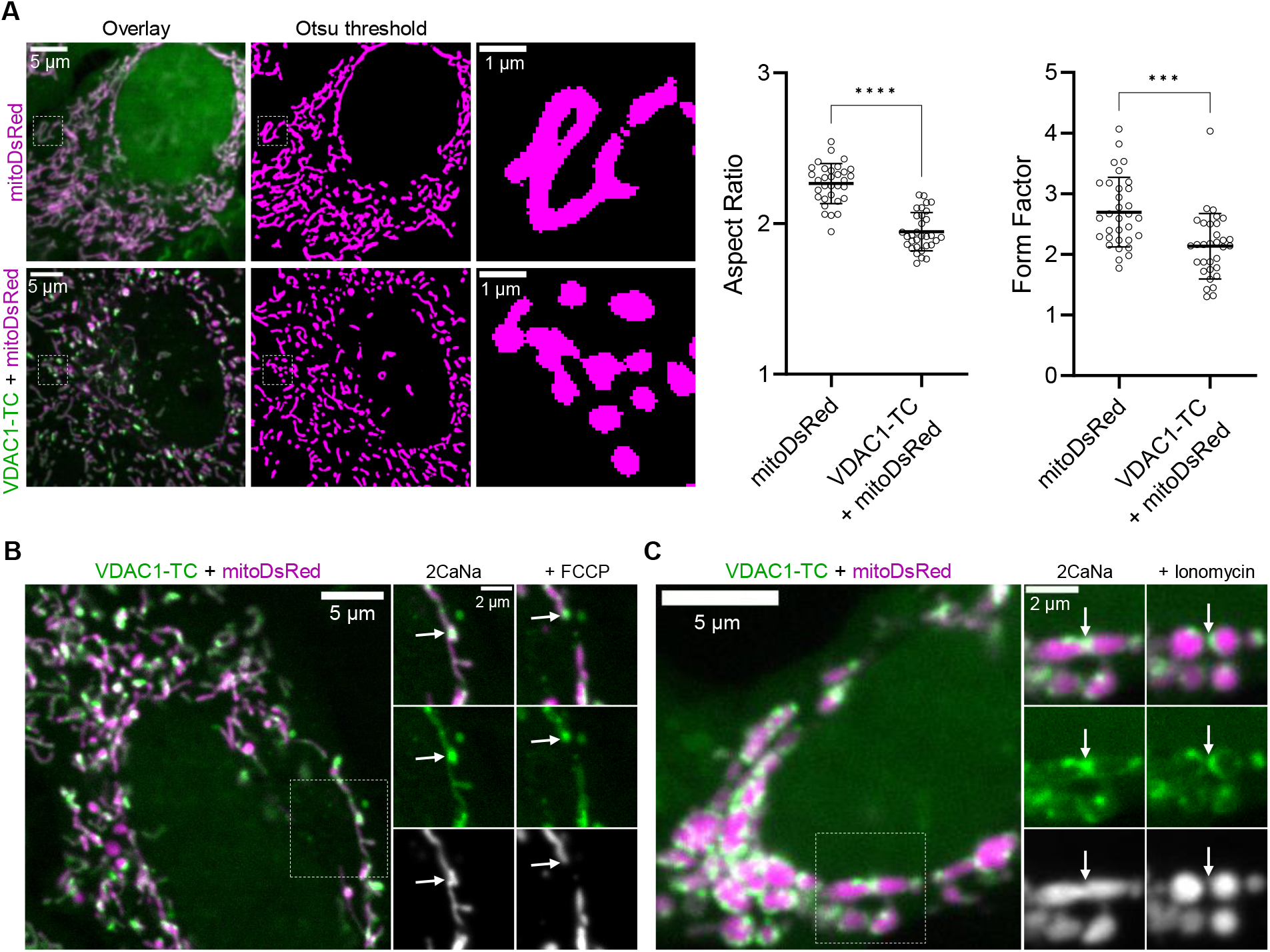
VDAC1 overexpression promotes mitochondrial fragmentation. (A) Confocal images of a HeLa cell expressing mitoDsRed (top row) or VDAC1-TC with mitoDsRed (bottom row); cells were labeled with 1 µM FlAsH for 15 min and washed with 100 µM BAL for 10 min. The left column shows an overview of the cell, the middle column shows the Otsu thresholded mitoDsRed signal, and the dashed squares are magnified in the right column. Graphs on the right show the Aspect Ratio and Form Factor of thresholded images. The difference between groups was evaluated using unpaired t-test. Data are presented as mean ± SD. (B) Confocal images of a HeLa cell expressing VDAC1-TC and mitoDsRed; the cell was labeled with 1 µM ReAsH for 15 min and washed with 100 µM BAL for 10 min (right). On the left, an overview of the cell is shown, and the dashed square is magnified on the right before (middle) and 10 min after perfusion with 2 µM FCCP. White arrows point to a VDAC1-cluster before and after mitochondrial fission. (C) Confocal images of a HeLa cell expressing VDAC1-TC and mitoDsRed; the cell was labeled with 1 µM ReAsH for 15 min and washed with 100 µM BAL for 10 min (right). On the left, an overview of the cell is shown, and the dashed square is magnified on the right before (middle) and 10 min after perfusion with 4 µM Ionomycin. White arrows point to a VDAC1-cluster before and after mitochondrial fission.

## Discussion

Here, we demonstrate the efficacy of the short TC-tag in visualizing the dynamics and interactions of VDAC1 in living cells. By overcoming the limitations associated with larger FPs, the TC-tag offers a valuable tool for studying the roles of VDAC1 in cellular processes, such as ER-mitochondrial communication, apoptosis, and mitochondrial dynamics.

Our data suggest that N- and C-terminal fusion of FPs leads to mistargeting and aggregation of VDAC1 in HeLa cells. This is in line with previous reports, showing poor targeting of C-terminal FP-tagged rat and human VDAC1 in HeLa cells.^3,4^ Using a short TC-tag, we could visualize VDAC1-clusters on mitochondria. This cluster-like distribution of VDAC1 has also been observed with FLAG-tagged VDAC1 and endogenously immunolabeled VDAC1.^14,15^ We demonstrated that VDAC1-clusters are localized at ER-mitochondria contact sites using high-resolution fluorescence microscopy. Since studies suggest that VDAC1 is localized at ER-mitochondrial contact sites,^11,12^ this finding further indicates that TC-tagged VDAC1 reflects the same localization patterns as endogenous VDAC1.

Notably, we discovered a frequent colocalization between VDAC1- and BAK-clusters during glucose depletion or STS-induced apoptosis. While VDAC1 has been suggested to be involved in BAK-mediated apoptosis,^13^ there was no evidence of their colocalization until now.

Furthermore, we observed that VDAC1 localized to mitochondrial fission sites when cells were treated with FCCP or ionomycin. This spatial correlation suggests that VDAC1 is involved in mitochondrial fission events. We also found that VDAC1 overexpression led to a marked increase in mitochondrial fragmentation. While VDAC1 overexpression could affect mitochondrial dynamics, our study is in line with a previous report demonstrating that the knockdown of VDAC1 prevents mitochondrial fragmentation induced by glutamate excitotoxicity in cultured neurons.^16^

In conclusion, we introduce a novel approach to tag VDAC1 and allow precise imaging of VDAC1 in live cells. We anticipate that this approach will help unravel the role of VDAC1 in the pathophysiology of living cells.

## Methods

### Cell culture

HeLa cells were cultured in Dulbecco’s modified Eagle’s medium (DMEM D5523, Sigma-Aldrich) supplemented with 10% FCS, 10 mM NaHCO_3_, 50 U/mL penicillin-streptomycin, 1.25 µg/mL amphotericin B and 25 mM HEPES; pH was adjusted to 7.45 with NaOH. Cells were grown in a humified atmosphere of 5% CO_2_ at 37°C.

### Transfection

Cells were seeded in 6-well plates on 30 mm glass coverslips (Paul Marienfeld GmbH & Co. KG, Lauda-Königshofen, Germany) and transfected using PolyJet (SignaGen Laboratories). Per well, 3 µl of PolyJet reagent was mixed with 1 µg of plasmid DNA in 100 µl of DMEM devoid of serum and antibiotics. The transfection mixture was added to 1 ml of culture medium for 8 hours and was then replaced with 2 ml of culture medium. Imaging was performed 24 - 48 h after transfection.

### Imaging of subcellular protein dynamics

Before imaging, cells were put into a storage buffer, which was composed of 135 mM NaCl, 5 mM KCl, 2 mM CaCl_2_, 1 mM MgCl_2_, 10 mM HEPES, 2.6 mM NaHCO_3_, 0.44 mM KH_2_PO_4_, 0.34 mM Na_2_HPO_4_, 10 mM D-glucose, 2 mM L-glutamine, 1X MEM amino, 1X MEM vitamins, 1% penicillin-streptomycin and 1% Amphotericin B; pH was adjusted to 7.45 with NaOH.

High-resolution imaging was performed with an array confocal laser scanning microscope (Axiovert 200 M, Zeiss) equipped with a 100×/1.45 NA oil immersion objective (Plan-Fluor, Zeiss) and a Nipkow-based confocal scanner unit (CSU-X1, Yokogawa Electric Corporation). Laser light of diode lasers (Visitron Systems, Pucheim, Germany) served as the excitation light source: CFP, GFP, and RFP fusion constructs were excited with 445, 488, and 561 nm lasers, respectively. Emission light was captured with a CoolSNAP HQ2 CCD Camera (Photometrics Tucson, Arizona, USA) using the emission filters ET460/50m, ET525/36m, and ET630/75m (Chroma Technology Corporation) for CFP, GFP, and RFP fusion constructs, respectively. FlAsH was excited like GFP for two-color imaging and like YFP for three-color imaging (excitation laser: 514 nm; emission filter ET535/30m). ReAsH was excited like RFP.

### Image analysis

Image analysis was performed with Fiji software. Z-stack images with a step size of 200 nm were deconvoluted and background-subtracted using a rolling ball radius of 50 to 300 pixels. For colocalization analysis, the TrackMate plugin was used to identify and quantify interactions between VDAC1-clusters and the ER or BAK-clusters. To assess mitochondrial morphology, 2D images of a mitochondrial matrix marker were thresholded using the Otsu method. For the Aspect Ratio, the ratio of the longest and shortest axes of a mitochondrial matrix marker was calculated. For the Form Factor, the following formula was used: 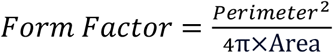

### VDAC1-TC labeling

Cells expressing tetracysteine-tagged VDAC1 (VDAC1-TC) were stained with FlAsH-EDT_2_ or ReAsH-EDT_2_ (Cayman chemical, Michigan USA) in EHL for 15 min at 37°C. Dye concentrations are indicated in figure legends. The cells were washed with 100 µM BAL (2,3-dimercaptopropanol or British anti-Lewisite) in EHL for 10 min at 37°C and kept in EHL before confocal imaging.

### Perfusion of cells during live cell imaging

Transfected cells on 30 mm coverslips were put in a PC30 perfusion chamber (NGFI, Graz, Austria) and perfused at a flow rate of approximately 1 ml per minute with a gravity-based perfusion system (PS9, NGFI). Glucose buffer was composed of 135 mM NaCl, 5 mM KCl, 2 mM CaCl_2_, 1 mM MgCl_2_, 10 mM HEPES, and 10 mM D-glucose; pH was adjusted to 7.45 with NaOH. The glucose-free buffer contained 10 mM D-mannitol instead of glucose.

### Chemicals

To induce mitochondrial fission, cells were perfused with 2 µM FCCP (Sigma) or 4 µM Ionomycin (Abcam) for 10 min in the glucose buffer. To induce apoptosis, cells were incubated with 10 µM staurosporine for one hour in EHL.

### Generation of 3D structures of VDAC1-fusion constructs

3D structures of GFP-VDAC1, VDAC1-GFP, and VDAC1-TC were predicted using ColabFold v1.5.2-patch, a modified version of AlphaFold2 that integrates MMseqs2 for sequence alignment and structure prediction. 3D structures were visualized using UCSF Chimera.

### Plasmids

| Name | Description | Reference |
| --- | --- | --- |
| GFP-VDAC1 | GFP fused to the N-terminus of VDAC1 | This study |
| VDAC1-VDAC1 | GFP fused to the C-terminus of VDAC1 | This study |
| VDAC1-TC | TC-tag fused to the C-terminus of VDAC1 | This study |
| mitoDsRed | Mitochondrial matrix marker | Addgene #44386 |
| HK1-GFP | HK1 marker | Addgene #21917 |
| mCh-ER3 | ER marker | Addgene #55041 |
| GFP-BAK | BAK marker | Addgene #32564 |

## Acknowledgments

We appreciate the technical assistance from René Rost, Anna Schreilechner, and Mercedes Maier. The research was supported by the Molecular Medicine PhD program of the Medical University of Graz and the FWF (Austrian Science Fund: I3716-B27 to R.M.). We are grateful to the reviewers and editors for their precious time.

## Author Contributions

J.P. and R.M. conceived the study. J.P. performed experiments and carried out image analysis. F.E.O., B.G., Y.C.E, W.F.G., and R.M. contributed to the interpretation of results and experimental design. J.P. and R.M. wrote the first draft of the manuscript, and all authors contributed to the final version.

---

**Figure S1.**
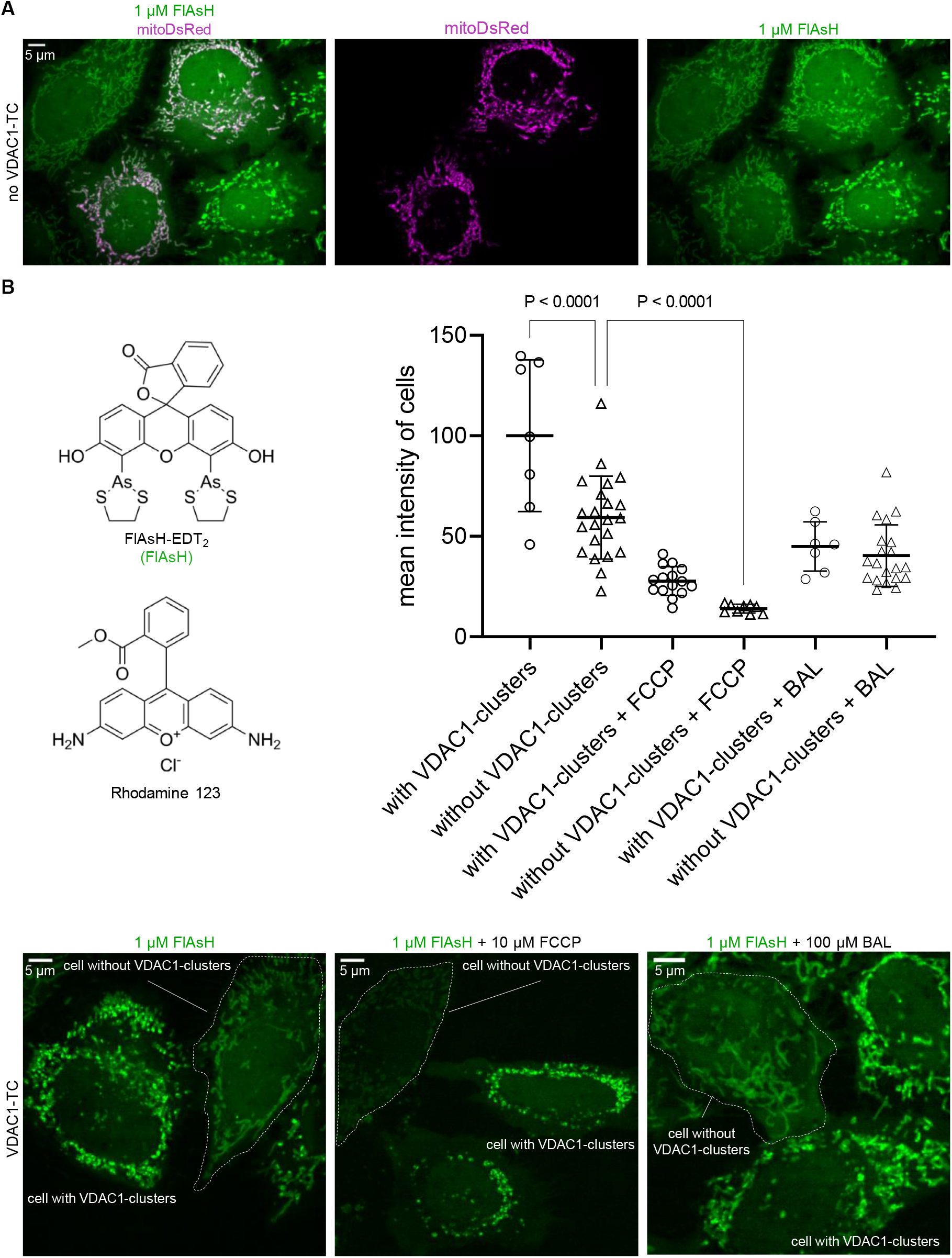
Short tetracysteine-tag allows to visualize VDAC1-clusters on mitochondria. (A) Confocal images of HeLa cells transfected with mitoDsRed; cells were labeled with 1 µM FlAsH for 15 min. (B) Chemical structures of FlAsH and Rhodamine123 (left). Confocal images of HeLa cells transfected with VDAC1-TC; cells were labeled with 1 µM FlAsH for 30 min, in the presence of 10 µM FCCP or washed with 100 µM BAL for 10 min (bottom). The dashed outline indicates cells without VDAC1-clusters. The mean intensity of cells with and without VDAC1-clusters was measured (right graph). The difference between groups was evaluated using one-way ANOVA with Bonferroni post hoc test. Data are presented as mean ± SD.

## References

1. Nienhaus, K., and Nienhaus, G.U. (2022). Genetically encodable fluorescent protein markers in advanced optical imaging. Methods Appl. Fluoresc. 10.

2. Kremers, G.-J., Gilbert, S.G., Cranfill, P.J., Davidson, M.W., and Piston, D.W. (2011). Fluorescent proteins at a glance. J. Cell Sci. 124, 157–160.

3. Rapizzi, E., Pinton, P., Szabadkai, G., Wieckowski, M.R., Vandecasteele, G., Baird, G., Tuft, R.A., Fogarty, K.E., and Rizzuto, R. (2002). Recombinant expression of the voltage-dependent anion channel enhances the transfer of Ca2+ microdomains to mitochondria. J. Cell Biol. 159, 613–624.

4. Dubey, A.K., Godbole, A., and Mathew, M.K. (2016). Regulation of VDAC trafficking modulates cell death. Cell death Discov. 2, 16085.

5. Kozjak-Pavlovic, V., Ross, K., Götz, M., Goosmann, C., and Rudel, T. (2010). A Tag at the Carboxy Terminus Prevents Membrane Integration of VDAC1 in Mammalian Mitochondria. J. Mol. Biol. 397, 219–232.

6. Jores, T., Klinger, A., Groß, L.E., Kawano, S., Flinner, N., Duchardt-Ferner, E., Wöhnert, J., Kalbacher, H., Endo, T., Schleiff, E., et al. (2016). Characterization of the targeting signal in mitochondrial β-barrel proteins. Nat. Commun. 7, 12036.

7. Griffin, B.A., Adams, S.R., and Tsien, R.Y. (1998). Specific covalent labeling of recombinant protein molecules inside live cells. Science 281, 269–272.

8. Hoffmann, C., Gaietta, G., Zürn, A., Adams, S.R., Terrillon, S., Ellisman, M.H., Tsien, R.Y., and Lohse, M.J. (2010). Fluorescent labeling of tetracysteine-tagged proteins in intact cells. Nat. Protoc. 5, 1666–1677.

9. Shoshan-Barmatz, V., Krelin, Y., Shteinfer-Kuzmine, A., and Arif, T. (2017). Voltage-Dependent Anion Channel 1 As an Emerging Drug Target for Novel AntiCancer Therapeutics. Front. Oncol. 7, 154.

10. Abu-Hamad, S., Zaid, H., Israelson, A., Nahon, E., and Shoshan-Barmatz, V. (2008). Hexokinase-I protection against apoptotic cell death is mediated via interaction with the voltage-dependent anion channel-1: mapping the site of binding. J. Biol. Chem. 283, 13482–13490.

11. Szabadkai, G., Bianchi, K., Várnai, P., De Stefani, D., Wieckowski, M.R., Cavagna, D., Nagy, A.I., Balla, T., and Rizzuto, R. (2006). Chaperone-mediated coupling of endoplasmic reticulum and mitochondrial Ca2+ channels. J. Cell Biol. 175, 901–911.

12. Liu, Y., Ma, X., Fujioka, H., Liu, J., Chen, S., and Zhu, X. (2019). DJ-1 regulates the integrity and function of ER-mitochondria association through interaction with IP3R3-Grp75-VDAC1. Proc. Natl. Acad. Sci. U. S. A. 116, 25322–25328.

13. Tajeddine, N., Galluzzi, L., Kepp, O., Hangen, E., Morselli, E., Senovilla, L., Araujo, N., Pinna, G., Larochette, N., Zamzami, N., et al. (2008). Hierarchical involvement of Bak, VDAC1 and Bax in cisplatin-induced cell death. Oncogene 27, 4221–4232.

14. Neumann, D., Bückers, J., Kastrup, L., Hell, S.W., and Jakobs, S. (2010). Two-color STED microscopy reveals different degrees of colocalization between hexokinase-I and the three human VDAC isoforms. PMC Biophys. 3, 4.

15. Durel, B., Kervrann, C., and Bertolin, G. (2021). Quantitative dSTORM superresolution microscopy localizes Aurora kinase A/AURKA in the mitochondrial matrix. Biol. cell 113, 458–473.

16. Oppermann, S., Mertins, B., Meissner, L., Krasel, C., Psakis, G., Reiß, P., Dolga, A.M., Plesnila, N., Bünemann, M., Essen, L.-O., et al. (2021). Interaction between BID and VDAC1 is required for mitochondrial demise and cell death in neurons. bioRxiv, 2021.09.14.460262.

