## Supplementary Information for "Visualizing interactions of VDAC1 in live cells using a tetracysteine tag"

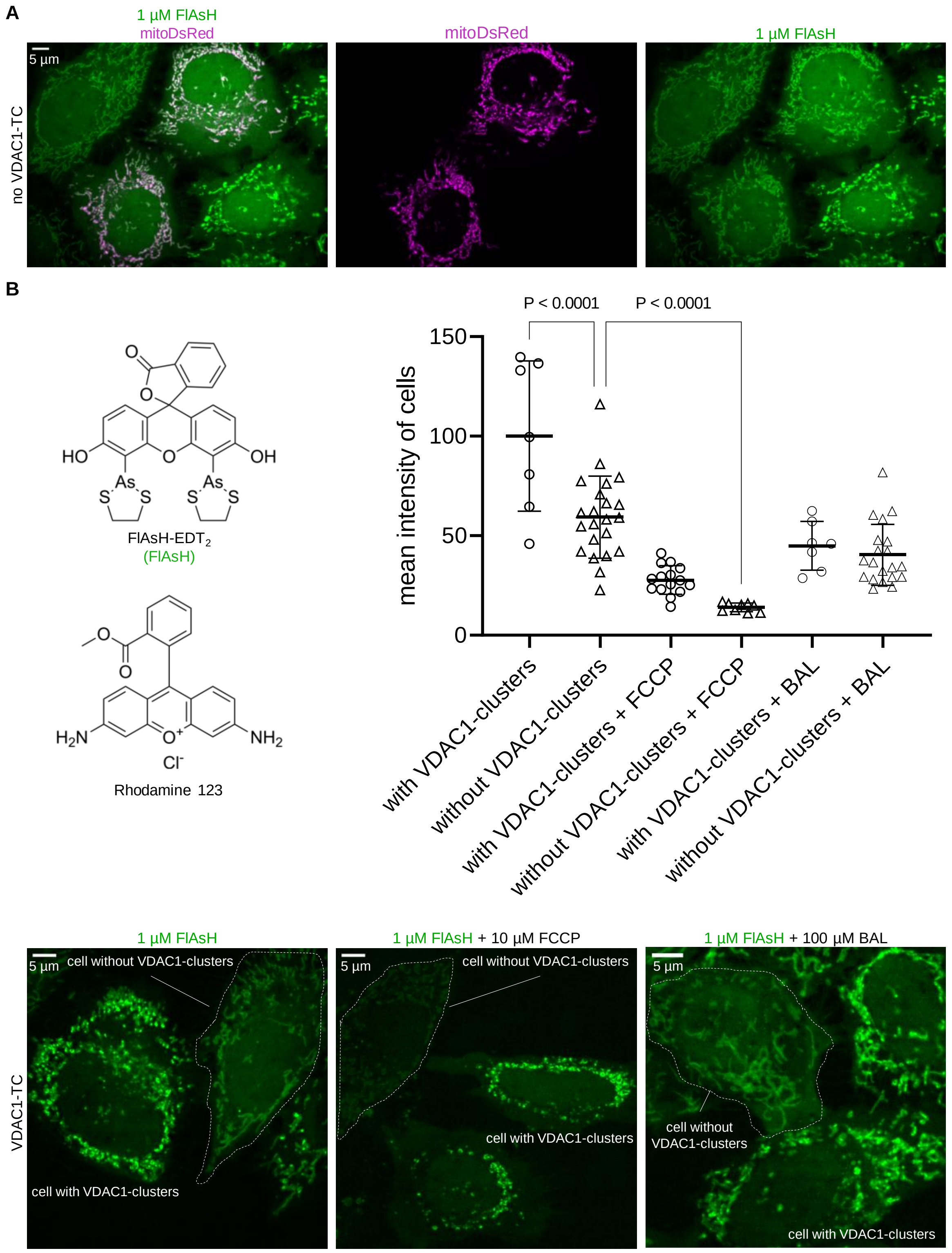
A
1 µM FlAsH
mitoDsRed
mitoDsRed
1 µM FlAsH
5 µm
no VDAC1-TC
B
HO
OH
As
S
S
As
S
S
O
O
O
FlAsH-EDT$_2$
(FlAsH)
O
O
$H_2N$
$O^+$
$NH_2$
$Cl^-$
Rhodamine 123
P < 0.0001
P < 0.0001
mean intensity of cells
150
100
50
0
with VDAC1-clusters
without VDAC1-clusters
with VDAC1-clusters + FCCP
without VDAC1-clusters + FCCP
with VDAC1-clusters + BAL
without VDAC1-clusters + BAL
1 µM FlAsH
1 µM FlAsH + 10 µM FCCP
1 µM FlAsH + 100 µM BAL
5 µm
cell without VDAC1-clusters
cell with VDAC1-clusters
VDAC1-TC
5 µm
cell without VDAC1-clusters
cell with VDAC1-clusters
5 µm
cell without
VDAC1-clusters
cell with VDAC1-clusters

**Figure S1. Short tetracysteine-tag allows to visualize VDAC1-clusters on mitochondria.** (A) Confocal images of HeLa cells transfected with mitoDsRed; cells were labeled with 1 µM FlAsH for 15 min. (B) Chemical structures of FlAsH and Rhodamine123 (left). Confocal images of HeLa cells transfected with VDAC1-TC; cells were labeled with 1 µM FlAsH for 30 min, in the presence of 10 µM FCCP or washed with 100 µM BAL for 10 min (bottom). The dashed outline indicates cells without VDAC1-clusters. The mean intensity of cells with and without VDAC1-clusters was measured (right graph). The difference between groups was evaluated using one-way ANOVA with Bonferroni post hoc test. Data are presented as mean ± SD.
